# Quantifying live bacterial densities using non-invasive optical measurements of *E. coli*

**DOI:** 10.1101/2021.06.12.448182

**Authors:** Eric van der Helm, Stephanie M. A. Redl

## Abstract

Profiling the growth of bacterial cultures over time can be a tedious and error-prone process. Here, we present the development and evaluation of the use of the ODity platform to optically measure bacterial cell densities non-invasively. The digital growth data for *E. coli* MG1655 was calibrated against colony forming units (CFU/mL) obtained by plating on solid media. Diauxic-like shifts of liquid *E. coli* MG1655 cultures grown at 37°C in LB media were observed at densities as low as 2.9 × 10^7^ ± 1.2 CFU/mL. The shift occurred at a significantly higher cell density (6.0 × 10^7^ ± 1.2 CFU/mL) when the bacteria were cultured at 31°C. These shifts were only short lived, 15.2 ± 1.5 and 20.8 ± 1.8 min at 37°C and 31°C, respectively, with the previous growth rate restored thereafter. We measured minimum doubling times of 17.0 ± 1.1 and 24.8 ± 0.9 min at 37°C and 31°C, respectively. These results demonstrate that the growth and growth rate of bacterial cultures can be accurately determined non-invasively using the ODity device.

## Introduction

Profiling the growth of a bacterial culture is tedious – the optical density of culture samples must be measured at regular intervals over a long period of time in order to construct each growth curve. Small-scale (e.g., 1 L) fermenters have been traditionally used to assess growth. However, their cost prohibits many labs from being equipped with a sufficient number of these devices. Various DIY-solutions have previously been developed to measure the growth of cultures, such as the turbidistat^1^ and morbidostat^2^ devices. A more comprehensive review of various methods is provided by Pilizota and Yang^3^. Besides the significant costs of fermenters, sampling can disturb the culture conditions. Alternative approaches that involve cultivation of microbes in microplate readers do not resemble the growth conditions of cultivation in higher volumes owing to their limitations associated with culture volume, temperature control, gas atmosphere, mixing, and evaporation of liquid. Therefore, we have developed and assessed the capability of the ODity platform, a non-invasive optical device (Figure 1a) designed to measure the light scattering of a growing culture and quantify cell densities in real time.

**Figure 1.**
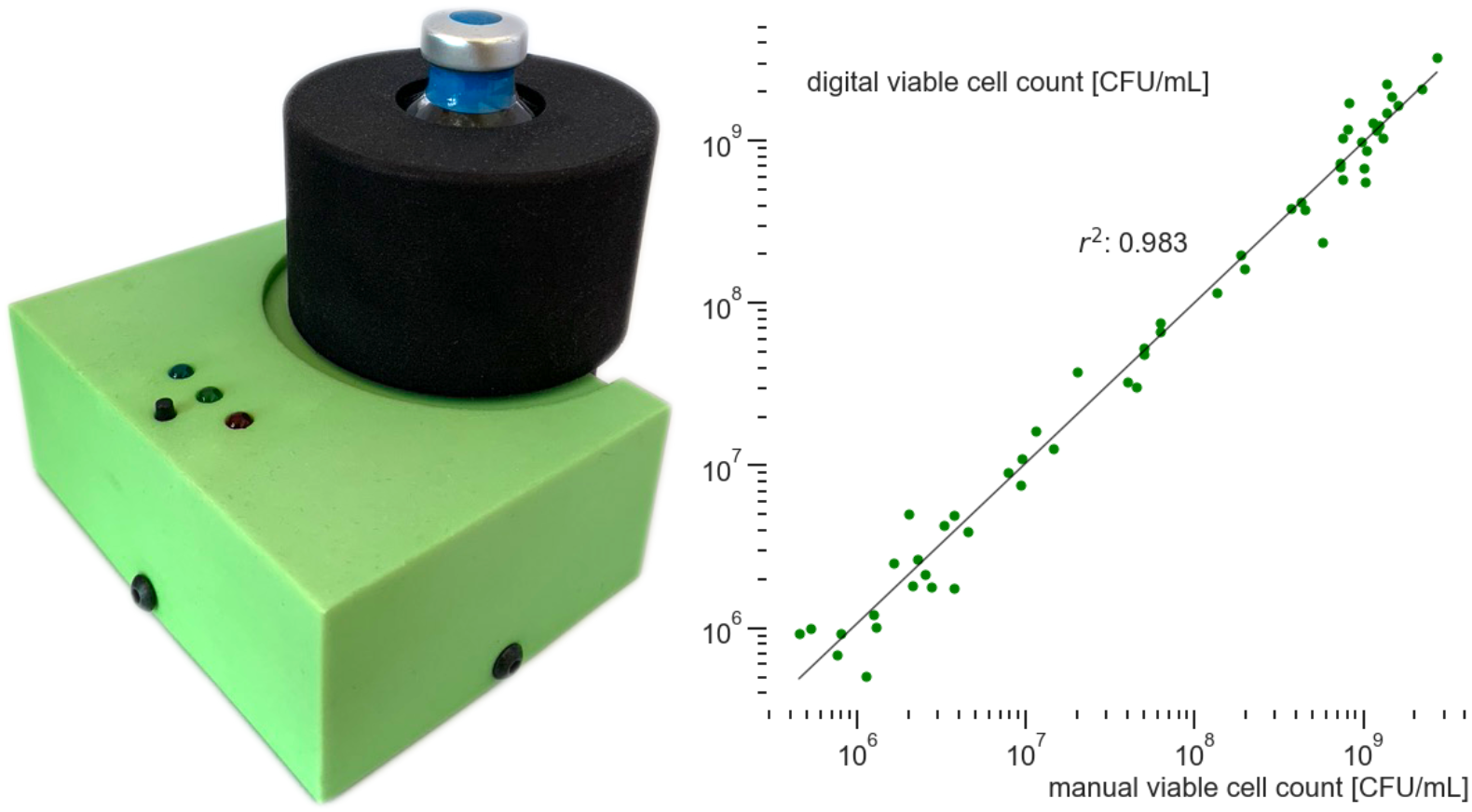
**(a)** Exterior of an ODity device D used to non-invasively measure the growth of a bacterial culture. **(b)** Relationship between the viable cell count as determined using counting of colony-forming units on solid media (horizontal axis) and measured using the ODity device (vertical axis). Complete linearity exists over a cell count data range across more than three orders of magnitude, with *r*^2^ = 0.983 and a detection limit approaching 10^6^ CFU/mL.

Although optical density (OD) is an often-used measure of cell growth, the metric has been difficult to interpret without calibration, owing to the influence of cell volume^4^ and cell size^5^ as well as its non-linear behavior^6^. Monod already noted^7^ this issue in 1949: *“[*..*]in spite of the widespread use of the optical techniques, not enough efforts have been made to check them against direct estimations of cell concentrations or bacterial densities”*. He even goes further in highlighting the importance of using temporally sampled measurements instead of only dilutions: *“Whatever instruments are used, the readings should be checked against bacterial density or cell concentration determinations, and the checks should be performed not only on different dilutions of a bacterial suspension, but at various times during the growth of a control culture. Only thus will the effects of variations of size of the cells be controlled. Without such controls it is impossible to decide whether the readings can be interpreted in terms of bacterial density or cell concentration, or both, or neither*.*”* Accordingly, we measured the light scattering of a growing *E. coli* MG1655 culture using the ODity device and enumerated the viable cells manually by plating them onto solid media. Viable cell counts are, in contrast to estimates based on optical density, independent of the aforementioned issues of cell size and non-linear behavior. We then used the ODity calibration software to establish a linear relationship between the obtained light scattering values against viable cells (colony-forming units, CFU) obtained from plating samples on solid medium (Figure 1b).

## Results and discussion

### E. coli growth curves

An *E. coli* MG1655 culture was inoculated into LB broth, starting at 10^3^ and 10^4^ CFU/mL and incubated at both 31°C and 37°C. The viable cell counts from nine replicates are summarized in Figure 2. The lag phase is below the limit of detection and thus not shown.

**Figure 2.**
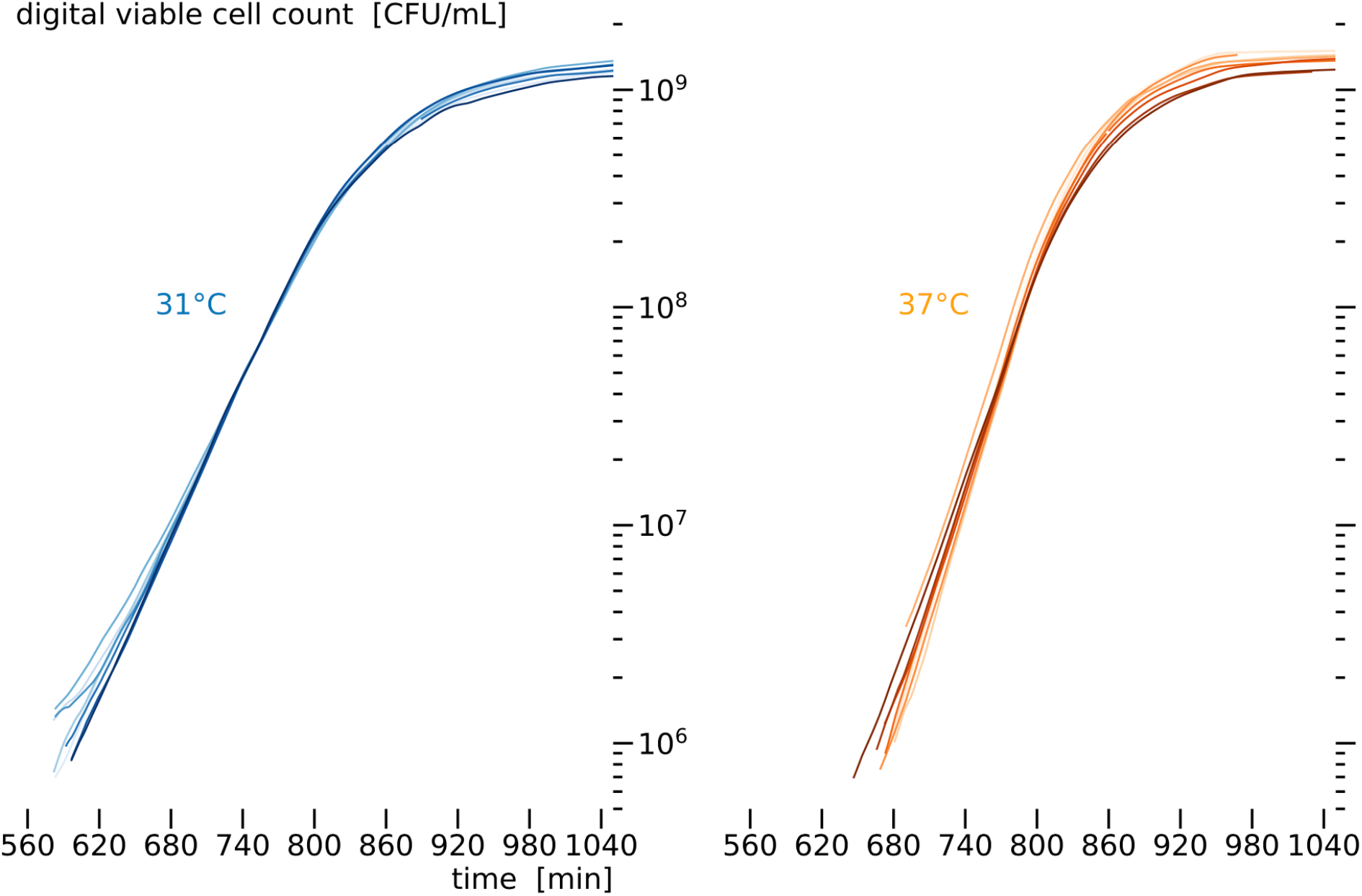
Examples of growth curves of *E. coli* MG1655 grown at 31°C (blue, *n* = 9) and 37°C (orange, *n* = 9) aligned in the time domain (horizontal axis).

As measurements were recorded at 2 min intervals, the growth rate (μ) over the whole measurement could be directly calculated without fitting growth models (Figure 3). The growth rate μ was calculated as 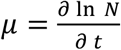 (Eq. 1), where ln *N* is the natural logarithm of the number of cells at a given time *t*.

**Figure 3.**
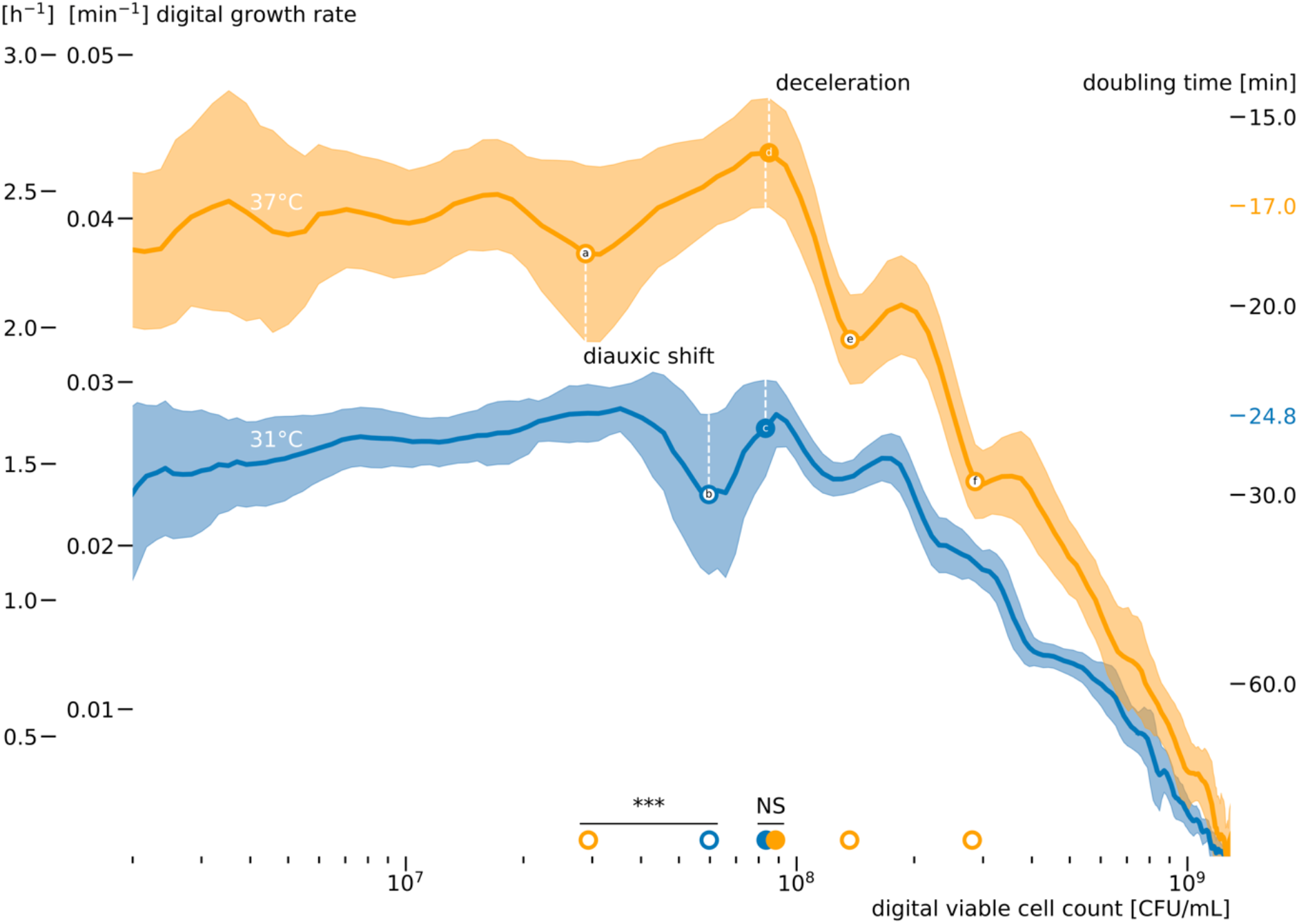
The digital growth rate of *E. coli* MG1655 cultured in LB broth at 37°C (orange) and 31°C (blue) as a function of the digital viable cell count measured using the ODity device. Shaded bands denote the standard deviation over nine biological replicates at 31°C (blue) and 37°C (orange); the growth data from Figure 2 are used throughout. When the cell density reaches the limit of detection at 10^6^ CFU/mL, the culture is already in the exponential phase. At 37°C, a temporary slowdown in growth was visible at 2.9 × 10^7^ ± 1.2 CFU/mL (marked with 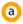) as previously reported by Sezonov and colleagues^9^ and attributed to a shift in carbon source utilization. At 8.8 × 10^7^ ± 1.1 CFU/mL, the exponential phase ended, and the deceleration phase began at 37°C (marked with 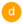), which was not significantly different (*p* = 0.11, Mann–Whitney U test) from the results for the 31°C cultures, at 8.4 × 10^7^ ± 1.1 CFU/mL (marked with 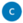). The deceleration phase was not uniform, and in both growth profiles, more distinct diauxic shifts can be recognized, with two more observable shifts for the 37°C culture at 1.4 × 10^8^ ± 1.1 (marked with 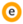) and 2.8 × 10^8^ ± 1.0 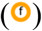 CFU/mL. In cultures maintained at 31°C, a similar diauxie was observed, albeit the minimum point was significantly (*p* = 2.1 × 10^−4^, Mann–Whitney U test) shifted upwards to 6.0 × 10^7^ ± 1.2 CFU/mL (marked with 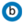). The minimum doubling times in the exponential phase at 37°C and 31°C were 17.0 ± 1.1 and 24.8 ± 0.9 min, respectively, as shown on the right axis. The layout of Figure 3 is analogous to that of Figure 2 published by Wang and Koch^8^.

### Diauxic shifts associated with culture temperature

In the culture grown at 37°C, at a density of 2.9 × 10^7^± 1.2 CFU/mL, a clear temporary drop in the growth rate was observed. This reduction in growth rate only lasted for 15.2 ± 1.5 and 20.8 ± 1.8 min at 37°C and 31°C, respectively (see Supplementary Figure 1), after which the previous growth rate was restored. The effect size (Hedges’ *g* = 3.25) of the duration difference between the cultures grown at 37°C and 31°C was significant (*p* = 5.2 × 10−^6^, Welch’s *t*-test).

This diauxic phenomenon was reported first by Wang and Koch^8^ in 1978 and later confirmed by Sezonov^9^ and colleagues in 2007. The latter researchers also offered a possible explanation: LB broth has a low concentration of sugars and thus mainly offers microbes amino acids as a carbon source. After depletion of the preferred carbon source at approximately 5 × 10^7^ CFU/mL (corresponding to OD = 0.3, as measured by ref ^9^), the cell metabolism switches to other carbon sources. However, Wang and Koch were unable to eliminate the pause in growth by supplementing LB broth with 19 L-amino acids, a trace element mixture, ZnCl_2_, or a mixture of adenosine, uridine, and cytidine. It should be noted that Wang and Koch utilized *E. coli* ML308 and not *E. coli* MG1655, as was used both by Sezonov and colleagues^9^ and in the present work.

In cultures grown at 31°C, a similar diauxie was observed, albeit the local minimum along the growth curve was significantly (*p* = 2.1 × 10^−4^, Mann–Whitney U test) shifted upwards to 6.0 × 10^7^ ± 1.2 CFU/mL, compared to 2.9 × 10^7^ ± 1.2 CFU/mL, at 37°C. Owing to the brief and transient nature of diauxie, such diauxic shifts often go unnoticed when measurement intervals exceed 5 min; furthermore, such shifts are not easily detected when measuring viable cells using plating (Supplementary Figure 2) and can be obscured by variation in standard laboratory procedures.

Although LB broth is easy to prepare, it is not a defined medium, and the composition can vary even between batches. Combined with the observations of diauxic growth relatively early in the growth trajectory, this inherent variability cautions against the use of LB broth when aiming to generate quantitative or physiological measurements reproducibly. Another interesting observation from Figure 3 is that the deceleration phase starts at the highest growth rate measured (a doubling time of 15 min). This could be owing to the previously mentioned diauxie, which acts as a temporary growth-synchronization event ^10^, however, more work is needed to elucidate this.

### Congruence of estimated growth rate with published estimates

From these experiments, the maximum growth rate was directly calculated with Eq. 2 and Eq. 3 based on nine biological replicates, providing an estimate of minimum doubling time of 17.0 ± 1.1 min at 37°C in LB broth.

This agrees with previous published reports (Table 1). When using the manually obtained viable cell counts by plating and fitting a four-parameter Baranyi–Roberts growth model^11^ (Supplementary Figure 2), we obtain a maximum growth rate of 0.035 ± 0.001 min^-1^, which is equivalent to a minimum doubling time of 19.8 min. Additionally, at 31°C, we obtain a maximum growth rate of 0.027 ± 0.001 min^-1^, which is equivalent to a minimum doubling time of 25.4 min. The Baranyi–Roberts growth model does not account for the temporary slowdown owing to diauxie and thereby slightly overestimates the minimum doubling time (and thus underestimates the maximum growth rate).

**Table 1.** Comparison of minimum doubling time of *E. coli* MG1655 cultured in LB at 37°C from selected references

| Minimum doubling time | Reference | Method | BIONumbers ID |
| --- | --- | --- | --- |
| 17.0 ± 1.1 | This study | Non-invasive high temporal resolution measurements using ODity |  |
| 18.1 ± 0.52 | Ref <sup>12</sup> | Time-lapse microscopy of 30 cells | 104331 |
| 19.8 | This study | Baranyi–Roberts model fit of viable cell count (CFU) |  |
| 20 | Ref <sup>9</sup> | Optical density (OD) | 103514 |
| 22.27 ± 0.69 | Ref <sup>12</sup> | Optical density (OD) |  |
| 25.7 | Ref <sup>4</sup> | Optical density (OD) calibrated by flow cytometry |  |

The minimum doubling time is defined as $t_{\text{minimum doubling time}} = \frac{\ln 2}{\mu_{\text{max}}}$ (Eq. 2), where the maximum growth rate is defined as $\mu_{\text{max}} = \max(\frac{\partial \ln N}{\partial t})$ (Eq. 3), with $\ln N$ being the natural logarithm of the number of cells at a given time $t$ .

## Conclusion

Here, we have used a non-invasive device to calibrate the measured light scattering of viable cells in a growing culture. As an example, we measure the growth of the model organism *E. coli* MG1655 in LB broth and found a diauxic growth minimum at 2.9 × 10^7^± 1.2 as reported previously^8,9^ at 37°C. We also showed this minimum shifts up to 6.0 × 10^7^ ± 1.2 when *E. coli* MG1655 is grown at 31°C. This reduction in growth rate only lasts for 15.2 ± 1.5 and 20.8 ± 1.8 min at 37°C and 31°C, respectively, after which the previous growth rate is restored. The short-lived nature of this shift may explain its scarce record in the published literature. The significant viable cell density at which the minimum of the diauxic shift occurs implies that the shift is temperature- and/or growth rate–dependent, and more experiments are needed to explain this phenomenon.

Furthermore, we estimated a minimum doubling time of 17.0 ± 1.1 min at 37°C in LB broth during the exponential phase and found that deceleration of the exponential phase already started at 8.8 × 10^7^± 1.1 CFU/mL.

Collectively, these results and their agreement with traditional methods and previously published results demonstrate that the growth and growth rate of bacterial cultures can be accurately determined non-invasively using the ODity device.

## Materials and methods

### Growth conditions

The starting inoculum was prepared from a single colony of streptomycin-resistant *E. coli* MG1655 (generated by spontaneous mutation as described in ref^13^) from an LB-agar plate. The colony was diluted in sterile PBS to a final concentration of 10^3^–10^4^ CFU/mL in 80 mL of sterile (pre-warmed) LB broth (with NaCl [5 g/L], tryptone [10 g/L], and yeast extract [5 g/L] in 1 L of sterile dH_2_O) supplemented with 150 μg/mL streptomycin.

The cultures were grown in clear glass serum bottles that were not capped to allow for air diffusion via a permeable membrane. The bottle contained a 25-mm Teflon-coated magnetic stir bar. The serum bottle was placed in an ODity Model D platform and secured in place with a screw cap. The unit was then placed onto a IKA RCT basic S001 magnetic stir plate at 350 rpm and moved into a semi-insulated enclosure to maintain a constant temperature.

### Measurements

The culture was illuminated at 624 nm, and scattering was measured at various angles. For the 18 experiments, a total of 6,274 measurements were recorded, separated by a median interval of 131.0 seconds.

The scattering was calibrated using the ODity cloud software against the viable cell count obtained by plating in colony forming units (CFU). Measurements were aligned in the time domain by minimizing the sum of distances of the first order derivative.

### Manual CFU measurements

To enumerate the bacterial concentration, 20 μL of the culture was 10-fold serially diluted in 180 μL of PBS and spotted onto LB-agar plates. Colony-forming units per milliliter of culture were calculated from the mean of two individually diluted replicates.

### Data processing

Data were manipulated using Pandas^14^, NumPy^15^, and SciPy^16^. The natural logarithm was used throughout this manuscript to calculate the growth rate. The growth data was filtered using a modified Savitzky–Golay function to account for non-uniformly spaced time points. A window size of 29 datapoints before 700 min or 7 datapoints after 700 min and a second order polynomial were used to compute the output. The Savitzky–Golay filter removes noise while maintaining the original signal structure, resulting in a more accurate derivative (in this case growth rate).

The growth rates of the individual biological replicates in Supplementary Figure 3 were merged by creating variable width bins with nine growth rate observations per cell density. The median of the nine cell densities per bin was assigned as the bin’s label. The mean and the standard deviation of the nine growth rates per bin were then used to construct Figure 3. The resulting data in Figure 3 was smoothed with a moving average (over five bins) to reduce noise. The glyphs marking the diauxic shift minima and deacceleration points were calculated as the mean from the individual biological replicates shown in Supplementary Figure 3. The maximum growth rate was calculated as an average of the growth rate over a period between the start of the diauxic shift and 40 min prior to the start of the diauxic shift.

### Baranyi–Roberts growth curves

Baranyi–Roberts growth curves were calculated as described by Baranyi and Roberts^11^ and, more specifically, according to their four parameter model^17^, in which the curvature parameters for deceleration and physiological adjustment rate are fixed, as follows:

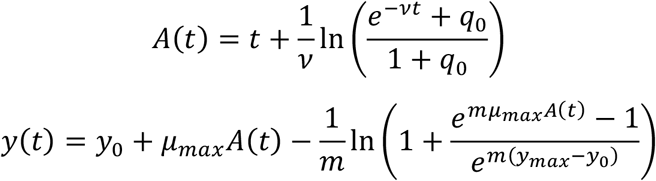

Fixing the curvature parameters *ν* = *μ*_*max*_ and *m* = 1 results in

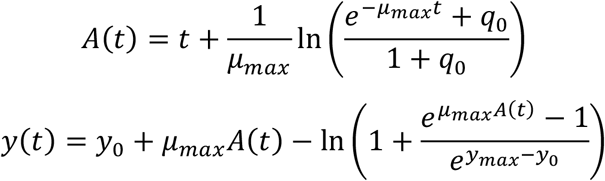

where y_0_ is the initial value of abundance, *μ*_*max*_ is the maximum growth rate (1/time), *y*_*max*_ is the carrying capacity (max. abundance), and *q*_0_ is a parameter specifying the initial physiological state of organisms (e.g., cells).

The fitting procedure was conducting using the Python package LM-FIT v 1.0.2^18^ using the Basin-hopping optimization method. The maximum growth rate or minimum doubling time reported was obtained by numerically calculating the growth rate of the resulting fit instead of using the μ_max_ parameter, as an analytical solution does not exist and the μ_max_ parameter can be inaccurate^19^.

### Statistical analysis

The geometric mean and geometric standard deviation (SD factor) are used throughout when calculating and reporting averages of manual and digital CFU quantities.

Welch’s *t*-test (as implemented in the SciPy function `ttest_ind` with `equal_var=False`) was used to compare the time duration distributions of the diauxic shift between the two different growth temperatures, as these distributions are assumed to be normally distributed but with potentially unequal variances. The effect size was calculated using Hedges’ *g*, which is more suitable for smaller sample sizes than Cohen’s *d*.

The minima of the diauxie for each growth rate profile were determined by searching for a local minimum in the exponential phase of the growth profile. Because the distributions are not assumed to be normal (for example, owing to the batch effect of the LB influencing the exact location of the local minima), the nonparametric Mann–Whitney *U* test (as implemented in the SciPy function `mannwhitneyu`) was used to compare the growth at 37°C and 31°C.

## Code availability

All the code used to generate the analysis and figures is publicly available in a Python Jupyter^20^ notebook on GitHub (https://github.com/EvdH0/ODity-growth-E-coli) and retrievable with the DOI http://doi.org/10.5281/zenodo.4937662. For convenience, a Docker container is provided to run the Jupyter notebook without the need to install the library dependencies.

## Supplementary figures

**Supplementary Figure 1.**
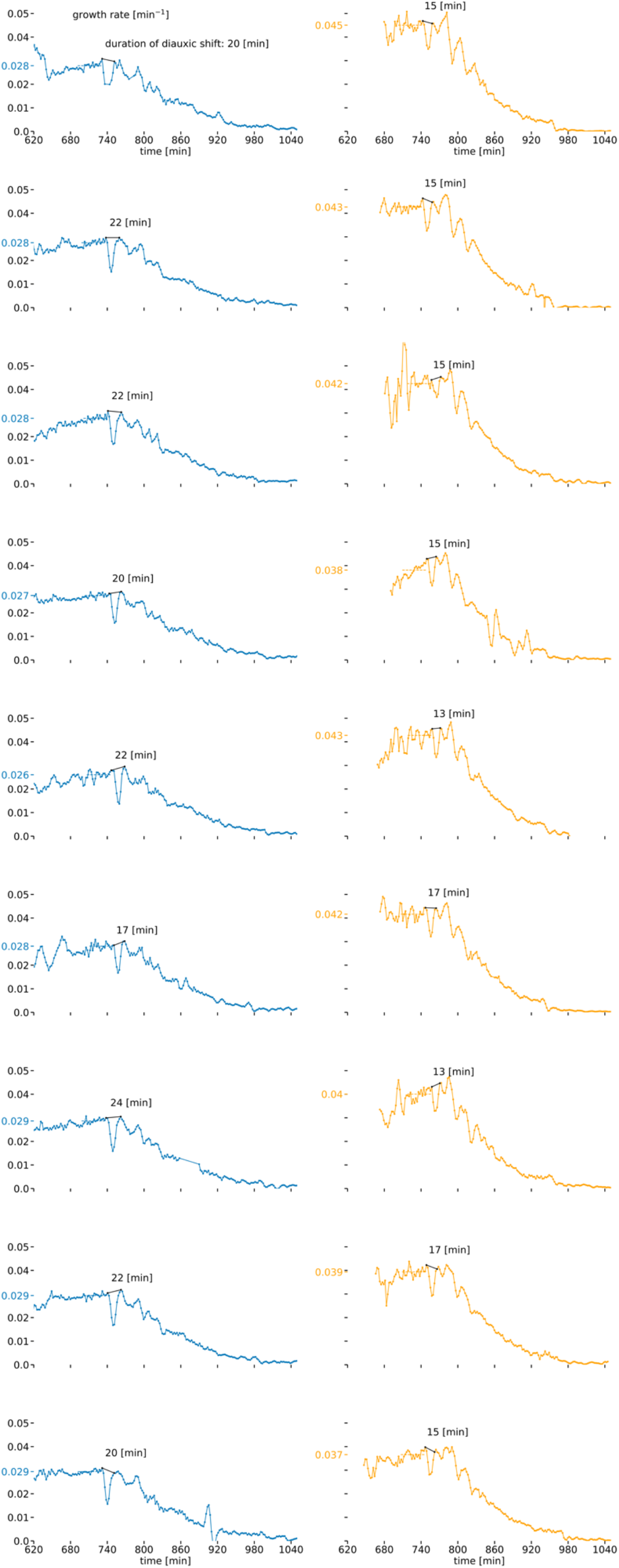
Growth rate (vertical) profiles of individual experiments of *E. coli* MG1655 grown on LB media at 31°C (left) and 37°C (right) showing the duration of the diauxic shift (indicated by the black solid line) and the determination of the growth rate (as an average over the period covered by the dashed line). The duration of the diauxic shift is calculated by identifying the local maxima around the previously identified local minimum and finally extracting the length shown in black. The maximum growth rate (tick mark on the axis) is calculated as the average of the growth rate over a period between the start of the diauxic shift and 40 min prior to the start of the diauxic shift. This reduction in growth rate only lasts for 15.2 ± 1.5 and 20.8 ± 1.8 min at 37°C and 31°C, respectively, after which the previous growth rate is restored. The effect size (Hedges’ *g* = 3.25) of the duration difference between the cultures grown at 37°C and 31°C was significant (*p* = 5.2 × 10^−6^, Welch’s *t*-test)

**Supplementary Figure 2.**
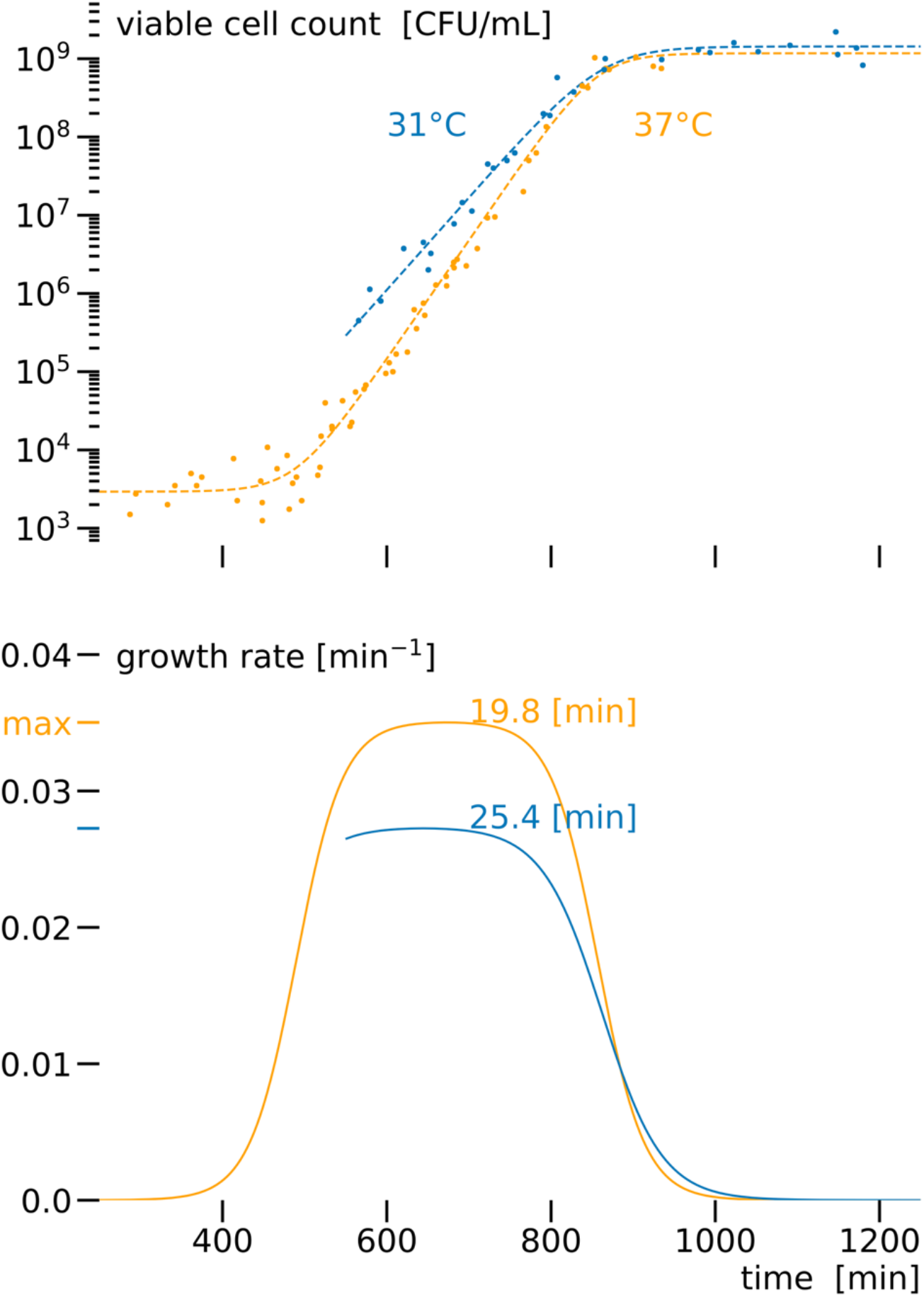
Estimation of growth rate based on fitting the Baranyi–Roberts growth model. Upper panel: manually measured viable cell count (CFU/mL) based on plating of aligned growth experiments at 31°C (blue) and 37°C (orange). Growth curves are estimated using the four parameter Baranyi–Roberts model^11,17^, shown as dashed lines. Lower panel: numerically calculated growth rate from the Baranyi–Roberts model fit, with an estimated minimum doubling time of 19.8 and 25.4 min at 37°C and 31°C, respectively.

**Supplementary Figure 3.**
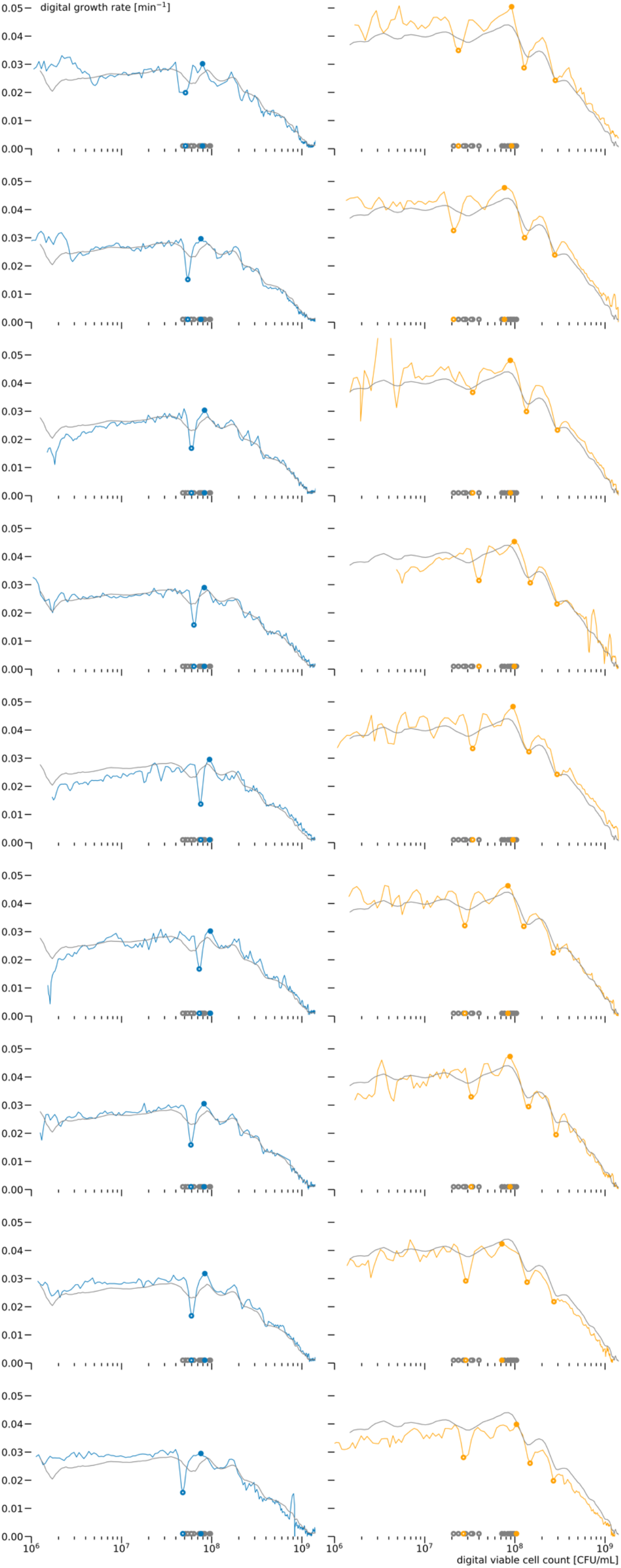
Growth rate (vertical) profiles of individual experiments of *E. coli* MG1655 grown on LB media at 31°C (left, blue) and 37°C (right, orange) as a function of digital viable cell count (CFU/mL) used for the determination of the locations of the diauxic shifts and deacceleration. The local minima associated with diauxic shifts are indicated by open markers 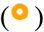, and the local maxima associated with deacceleration are indicated by solid markers 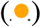. The gray bottom markers are the result of the other biological replicates at either 31°C or 37°C. The gray line denotes the average for either condition and corresponds to the average blue or orange line in Figure 3 in the main text.

